# A high-content microscopy drug screening platform for regulators of the extracellular digestion of lipoprotein aggregates by macrophages

**DOI:** 10.1101/2024.09.26.615160

**Authors:** Cheng-I J. Ma, Noah Steinfeld, Weixiang A. Wang, Frederick R. Maxfield

## Abstract

The recruitment of macrophages to the intima of arteries is a critical event in atherosclerotic progression. These macrophages accumulate excessive lipid droplets and become “foam cells”, a hallmark of atherosclerosis. Most studies focus on lipid accumulation through macrophage interaction with modified monomeric low-density lipoprotein (LDL). However, in the intima, macrophages predominantly encounter aggregated LDL (agLDL), an interaction that has been studied far less. Macrophages digest agLDL and generate free cholesterol in an extracellular, acidic, hydrolytic compartment. They form a tight seal around agLDL through actin polymerization and deliver lysosomal contents into this space in a process termed digestive exophagy. There is some evidence that inhibiting digestive exophagy to slow cholesterol accumulation in macrophages protects them from becoming foam cells and slows the progression of atherosclerotic lesions. Thus, understanding the mechanisms of digestive exophagy is critical. Here, we describe a high-content microscopy screen on a library of repurposed drugs for compounds that inhibit lysosome exocytosis during digestive exophagy. We identified many hit compounds and further characterized the effects that five of these compounds have on various aspects of digestive exophagy. In addition, three of the five compounds do not inhibit oxidized LDL-induced foam cell formation, indicating the two pathways to foam cell formation can be targeted independently. We demonstrate that this high-content screening platform has great potential as a drug discovery tool with the ability to effectively and efficiently screen for modulators of digestive exophagy.

## Introduction

Despite major advances in our understanding of the mechanisms that underlie cardiovascular diseases associated with atherosclerosis,^1^ these diseases are the leading cause of death worldwide.^2-4^ Macrophages in the arterial intima are a major contributor to atherosclerosis.^5-7^ Dysfunction of the vascular endothelium, often caused by the retention of low-density lipoprotein (LDL), leads to the recruitment of monocytes to the intima where they differentiate into macrophages.^7^ Disrupting macrophage recruitment to the intima by knocking out a macrophage chemokine receptor that detects chemokines secreted by endothelial cells has been shown to ameliorate atherosclerosis.^8-10^

One way that macrophages contribute to atherosclerotic plaque formation is through the accumulation of excessive lipid droplets filled with cholesteryl esters.^11-14^ These macrophage “foam cells” eventually die and are phagocytosed by other macrophages,^15^ which then also become overloaded with cholesteryl esters. For many years, the field has focused primarily on the role macrophage interactions with soluble cholesterol species, such as modified monomeric LDL (e.g., oxidized LDL), play in foam cell formation.^16^ Various receptors interact with these soluble lipoproteins, leading to endocytosis. However, in atherosclerotic lesions, the cholesterol species encountered by macrophages are predominantly aggregated LDL (agLDL) that is tightly linked to the extracellular matrix.^17-22^ Because agLDL is tightly linked to the extracellular matrix, aggregated lipoproteins cannot undergo receptor-mediated endocytosis or phagocytosis without first being released from the extracellular matrix. Thus, it is surprising that these lipoproteins are hydrolyzed by a lysosomal enzyme, lysosomal acid lipase (LAL).^23,24^

Our laboratory has described the formation of an extracellular, acidic, hydrolytic compartment (lysosomal synapse) by which macrophages interact with and hydrolyze agLDL in a process we have termed digestive exophagy.^23,25-33^ Following exocytosis of lysosomes, lysosomal synapses are acidified by V-ATPase in the plasma membrane, allowing LAL to hydrolyze cholesterol esters.^23^ These extracellular compartments are surrounded by F-actin rich membranes, which form complex invaginations that tightly seal the lysosomal synapse.^26,33^ Fragments of agLDL are released and can be endocytosed and delivered to late endosomes and lysosomes before being stored as cholesteryl esters in lipid droplets.^23,31^ Our laboratory has partially characterized the signal transduction processes that regulate various stages of digestive exophagy.^29,34^

It is possible that partially inhibiting digestive exophagy to slow the delivery of cholesterol to the macrophages would protect macrophages from becoming foam cells and slow the progression of atherosclerotic lesions. Thus, identifying modulators of digestive exophagy is critical, not only for the identification of disease-modifying treatments, but also for developing our understanding of the mechanisms of digestive exophagy. In this paper, to identify novel modulators of digestive exophagy, we developed and performed a high-content microscopy screen on a more than 2,000 compound library of repurposed drugs for compounds that inhibit lysosome exocytosis during digestive exophagy with the goal of identifying a novel use for approved or investigational drugs in the treatment of atherosclerosis. We identified many hit compounds and further characterized the effects that five of these compounds have on various aspects of digestive exophagy.

## Results

### Developing a high-content screen for modulators of lysosome exocytosis during digestive exophagy

Our laboratory uses a quantitative fluorescence microscopy assay to measure the exocytosis of lysosomes towards agLDL during digestive exophagy.^27,35^ In this experiment, we deliver biotin-FITC-dextran to macrophage lysosomes by overnight pinocytic uptake followed by a 3-hour chase. We then incubate Streptavidin-Alexa647-agLDL with macrophages for 90 minutes. In this time, biotin-FITC-dextran exocytosed towards Streptavidin-Alexa647-agLDL will bind tightly to the agLDL. Cells are then incubated with excess biotin in medium to bind any unoccupied streptavidin sites, fixed, and permeabilized with 1.3% Triton X-100 to remove unbound lysosomal biotin-FITC-dextran (Figure 1A). The biotin-FITC-dextran signal that colocalizes with Streptavidin-Alexa647-agLDL is quantified by confocal microscopy.

**Figure 1:**
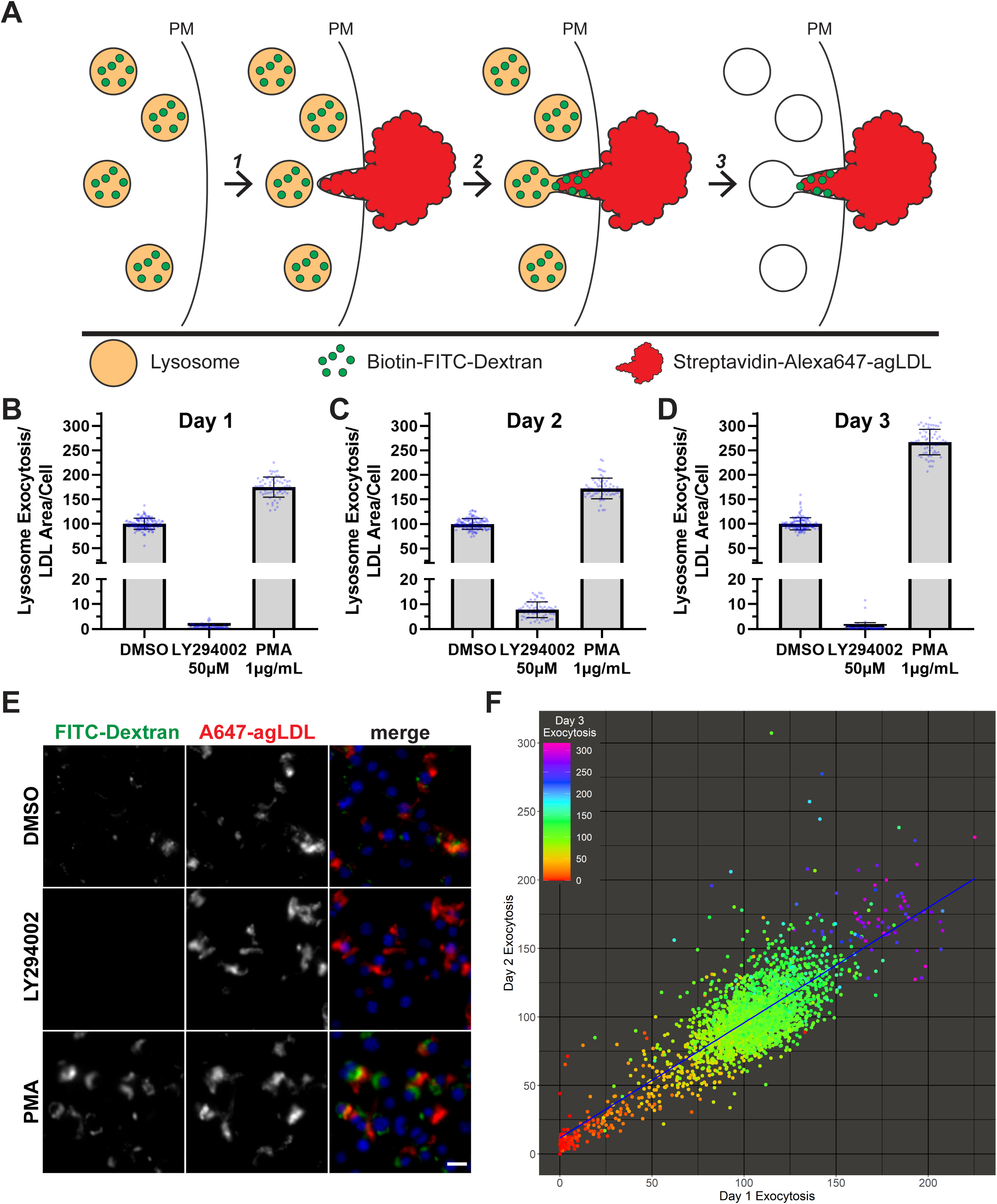
Developing a high-content screen for regulators of digestive exophagy. (**A**) Schematic of the lysosome exocytosis assay. Biotin-FITC-dextran is delivered to lysosomes by overnight pinocytic uptake followed by a 3-hour chase in media after washout. (1) Streptavidin-Alexa647-agLDL is added on top of J774 macrophages for 90 minutes to stimulate lysosome exocytosis. (2) After lysosome exocytosis, biotin-FITC-dextran binds tightly to Streptavidin-Alexa647-agLDL. Excess biotin is used to block any free streptavidin before fixation with 0.5% paraformaldehyde. (3) Permeabilization of the cells in PBS with 1.3% Triton X-100 is used to remove the unbound intracellular biotin-FITC-dextran before microscopy quantification. (**B-D**) On each of the eight 384-well plate used in the screen, column 23 contained DMSO-treated control cells (n=128) while half of column 24 contained cells treated with 50μM LY294002 (PI3K inhibitor) (n=64) and the other half of column 24 contained cells treated with 1μg/mL PMA (PKC activator) (n=64). J774 macrophages were assayed for lysosome exocytosis and normalized to both the agLDL area and the cell count. In each well, this measurement is expressed as a percentage of the average of the DMSO controls on its corresponding plate. Error bars indicate standard deviation. On Day 1 (**B**) the Z’ for LY294002 was 0.630 and the Z’ for PMA was -0.281. On Day 2 (**C**) the Z’ for LY294002 was 0.535 and the Z’ for PMA was -0.327. On Day 3 (**D**) the Z’ for LY294002 was 0.568 and the Z’ for PMA was 0.305. (**E**) Representative fluorescence micrographs from the screen, where DMSO (top), LY294002 (middle) and PMA (bottom)-treated J774 macrophages were assayed for lysosome exocytosis. Scale bar = 20μm. (**F**) Results from Day 1 of the screen were plotted against results from Day 2 of the screen with results from Day 3 of the screen indicated by color. A line of best fit is included in blue.

We adapted this assay for use in a high-content 384-well plate format. Compounds from the library of repurposed drugs were added to 384-well plates using a Janus Automated Workstation (Perkin-Elmer). J774 macrophages with biotin-FITC-dextran pre-internalized were plated on top of the compounds using a MultiDrop384 Dispenser (Thermo Fisher Scientific) and allowed to settle for 3 hours before adding Streptavidin-Alexa647-agLDL for 90 minutes. Subsequent PBS washes were performed using an ELx405 Microplate Washer (Bio-Tek Instruments), and excess biotin in medium, paraformaldehyde fixative, and Triton X-100 were added to each well sequentially using the MultiDrop384 Dispenser. The next day, 4 images in each well were acquired with an ImageXpress MICRO Automated High-Content Imaging System (Molecular Devices) using a 20x dry objective. This protocol is described in detail in the Materials and Methods section and was repeated on 3 days.

As a control to ensure that this assay was robust enough for a high-content format, on each 384-well plate, column 23 contained DMSO controls, and column 24 was split between the PI3-kinase inhibitor LY294002 (low signal control)^27^ and the protein kinase C activator phorbol 12-myristate 13-acetate (PMA, high signal control).^36^ These controls allow us to calculate the Z’ of the assay. Z’ is a metric defined by the means and standard deviations of positive and negative controls that attempts to quantify the suitability of an assay for use in a high-content screen.^37^ A Z’ between 0.5 and 1 is considered excellent, while a Z’ between 0 and 0.5 is considered somewhat marginal. For all 3 repeats of the screen, the Z’ for LY294002 was above 0.5, indicating the assay is suitable for a high-content format (Figure 1B-E). Surprisingly, the Z’ for PMA was less than 0 on days 1 and 2 and was only 0.305 on day 3, primarily because the magnitude of the effect of PMA on lysosome exocytosis during digestive exophagy was smaller in this screen than had been previously reported.^36^ This difference may be due to changes in the length of time that J774 cells were incubated with PMA before adding agLDL or in the way lysosome exocytosis is imaged and quantified. Consequently, hit compounds that decreased lysosome exocytosis are more likely to be *bona fide* hits than hit compounds that increased lysosome exocytosis.

Crucially, day to day screen results were highly reproducible (Figure 1F). Pairwise comparisons of each repeat of the screen each had an R^2^ correlation coefficient of nearly 0.7 (Supplementary Figure 1A-C). Furthermore, compounds with strongly decreased lysosome exocytosis on one day also tended to have strongly decreased lysosome exocytosis the other days. Together, these results demonstrate that the 384-well plate lysosome exocytosis assay was a robust tool for identifying novel regulators of digestive exophagy. Complete results from the screen are reported in Supplementary Table 1. In each well, lysosome exocytosis is normalized to both the agLDL area and the cell count, which is then expressed as a percentage of the average of the DMSO controls on its corresponding plate.

### Validation of five compounds that lowered lysosome exocytosis

We selected five hit compounds from the screen to validate. Three of these compounds had a more than 95% decrease in lysosome exocytosis compared to the DMSO control and are known to target proteins required for digestive exophagy. Lanraplenib inhibits spleen tyrosine kinase (SYK),^38^ and AZD-8186 is a potent and selective inhibitor of PI3Kβ and PI3Kδ.^39^ Our lab has previously implicated both of these kinases in digestive exophagy.^27^ LXS-196 (darovasertib) is a PKCα/θ inhibitor^40^ that also has a role in digestive exophagy.^36^ The fourth compound we validated, saracatinib,^41^ inhibits the protein tyrosine kinase SRC and was the strongest hit from the screen that did not kill a substantial number of cells. The final validated compound, lasofoxifene,^42^ is an estrogen receptor modulator that was developed to treat osteoporosis in postmenopausal women.^43^ While lasofoxifene only decreased lysosome exocytosis by around 65% in the screen, the parallels between digestive exophagy and the extracellular digestion of bone by osteoclasts during bone remodeling made it an interesting target for further investigation.^29^

To ensure that the results from the screen reflect general properties of macrophages, we tested whether the decrease in lysosome exocytosis observed for these five compounds in the screen in J774 immortalized macrophages could be recapitulated in bone marrow-derived macrophages (BMMs). To generate a dose response curve, we tested the effect varying concentrations of these five drugs had on lysosome exocytosis in BMMs. We tested concentrations from 10μM to 41nM in a series of 3-fold dilutions (Figure 2A-E). All five drugs effectively inhibited lysosome exocytosis at the highest concentration tested. Lanraplenib, AZD-8186, and saracatinib also exhibited more than 80% inhibition of lysosome exocytosis at 3.33μM while LXS-196 inhibited lysosome exocytosis by around 75% and lasofoxifene exhibited closer to 50% lysosome exocytosis reduction. These results closely resembled the results from the screen, which was performed with all drugs at a 5μM concentration. Having confirmed that these five compounds affect lysosome exocytosis in multiple types of macrophages, we performed further tests on the effects these compounds have on various aspects of digestive exophagy.

**Figure 2:**
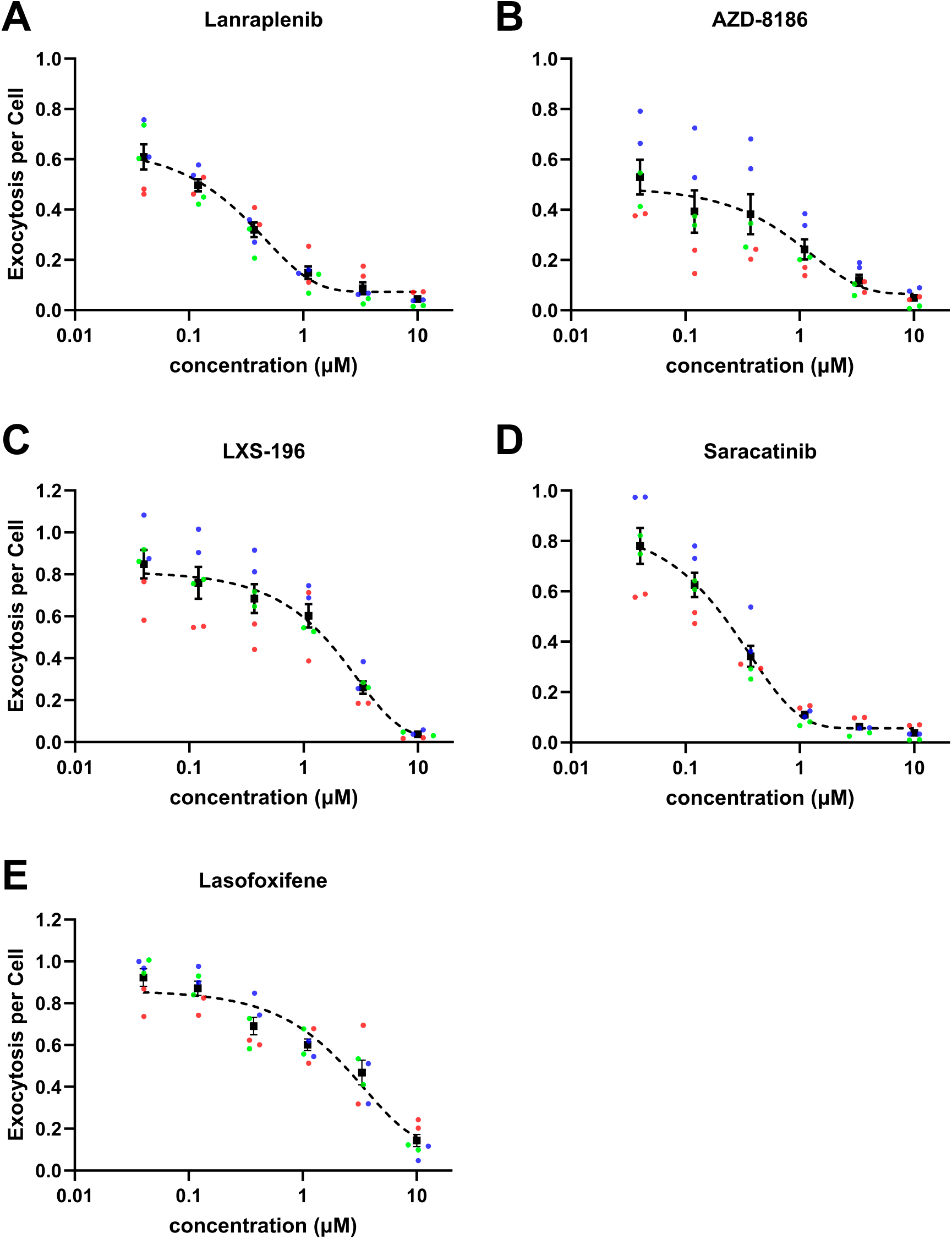
Determining the dose response curve of 5 hits from the screen. (**A-E**) BMMs were treated with DMSO or the indicated drug from 10μM to 41nM and assayed for lysosome exocytosis in a 96-well plate. Each black square represents an average of three independent experiments. Dashed line represents nonlinear regression curve fit of the data. (**A**) R^2^ = 0.9032 (**B**) R^2^ = 0.5904 (**C**) R^2^ = 0.8283 (**D**) R^2^ = 0.9091 (**E**) R^2^ = 0.8432.

### Each of the five validated compounds affects actin polymerization during digestive exophagy

In addition to lysosome exocytosis, the polymerization of filamentous actin (F-actin) around invaginations containing agLDL to generate a tightly sealed compartment is another critical step in digestive exophagy.^26,33,44^ To quantify the cell wrapping tightly around extracellular agLDL, after incubating fluorescent agLDL with macrophages for 1 hour, we stain F-actin with phalloidin and measure the colocalization of phalloidin with the agLDL as well as the total F-actin staining. Though F-actin does not truly colocalize with agLDL as it is localized just below the plasma membrane, by confocal microscopy using a 20x objective, agLDL and phalloidin appear effectively colocalized.

We examined the effect each of our five validated compounds had on actin polymerization during digestive exophagy in BMMs. The concentration of each drug was selected based on the dose response curve to be the lowest concentration at which we thought 80% reduction of lysosome exocytosis during digestive exophagy would be achieved. At the first concentration tested, lanraplenib (1μM) and AZD-8186 (500nM) had statistically significant, but small decreases in actin polymerization around agLDL (28% and 22%, respectively), with little or no effect on total F-actin levels (Supplementary Figure 2A-C). However, for lanraplenib especially, the results of this experiment were not consistent from day to day, suggesting that we were using a drug concentration near the IC50. Thus, we also tested the effect lanraplenib and AZD-8186 had on actin polymerization at double the concentration of the previous experiment (Figure 3A-C). At 2μM, lanraplenib had a much stronger effect on actin polymerization around agLDL, exhibiting a 66% decrease. AZD-8186 was only slightly more effective, exhibiting a 35% decrease in actin polymerization at 1μM. Neither drug at the higher concentration affected total F-actin levels. For lanraplenib, it is interesting that while the drug at 1.11μM has a more than 80% effect on lysosome exocytosis, at 1μM its effects on actin polymerization are inconsistent, perhaps suggesting that actin polymerization and lysosome exocytosis are inhibited at different concentrations for this drug.

**Figure 3:**
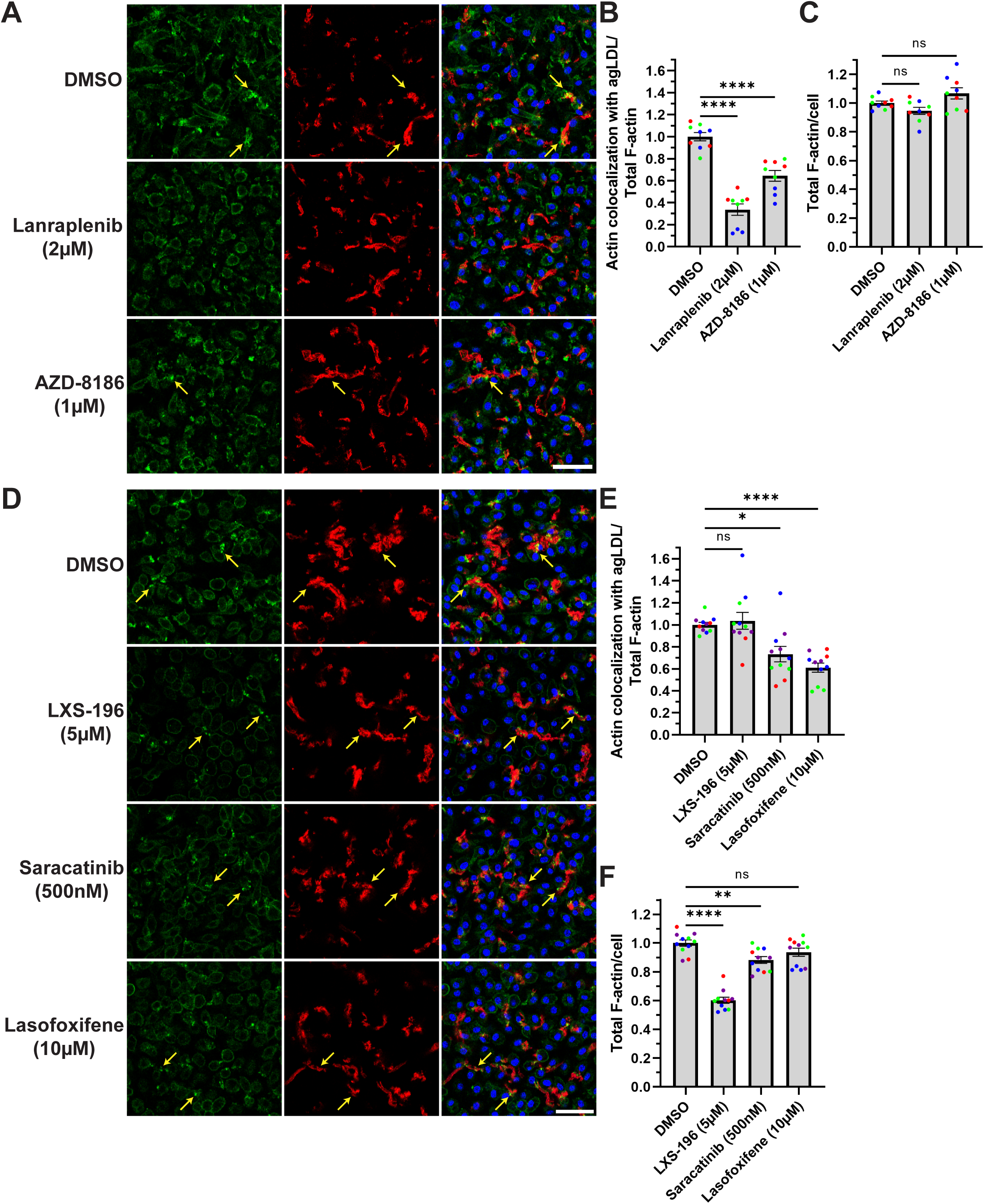
Actin polymerization around agLDL during digestive exophagy is affected by hits from the screen. (**A, D**) BMMs were treated with DMSO or the indicated drug and assayed for actin polymerization following a 1-hour incubation with Alexa647-agLDL in a 96-well plate. Scale bar = 50μm. Arrows highlight some areas of actin polymerization around agLDL. (**B, E**) Quantification of the colocalization of phalloidin-stained F-actin with agLDL normalized to total F-actin. (**C, F**) Quantification of total F-actin staining per cell.

We also examined the effect LXS-196, saracatinib, and lasofoxifene had on actin polymerization during digestive exophagy (Figure 3D-F). Treatment with LXS-196 did not result in less actin polymerization around agLDL, but did decrease total F-actin throughout the cell. This result is consistent with our previous report on the effect of LXS-196 on actin polymerization during digestive exophagy in J774 macrophages.^36^ Saracatinib and lasofoxifene exhibited modest decreases in actin polymerization around agLDL after normalizing for the changes in total F-actin. Saracatinib also exhibited a small but reproducible decrease in total F-actin levels.

Together, in addition to their strong effect on lysosome exocytosis during digestive exophagy, each of the five validated compounds also affect actin polymerization, though the magnitude of this effect varies among compounds.

### Testing whether the five validated compounds affect foam cell formation during digestive exophagy

To quantify the accumulation of lipids in the macrophages treated with agLDL, we measured the conversion of cholesterol from the cholesteryl esters in the agLDL into lipid droplets in macrophages.^28^ Following a 6 hour treatment with agLDL, cells are fixed and stained with LipidSpot 488, which stains neutral lipids (both agLDL and lipid droplets). We quantified intracellular lipid droplet formation as the LipidSpot 488 signal that does not colocalize with agLDL.

We examined the effect each of our five validated compounds had on lipid droplet accumulation during digestive exophagy in BMMs (Figure 4A, B). Apart from lasofoxifene, treatment with each of the compounds results in reduced accumulation of lipid droplets compared to a DMSO control. Interestingly, the same concentration of lanraplenib that exhibited inconsistent effects on actin polymerization effectively and consistently decreased lipid droplet accumulation. Most surprising, however, was the observation that treatment with lasofoxifene did not affect the accumulation of lipid droplets in cells treated with agLDL. It may be that in addition to inhibiting digestive exophagy, lasofoxifene accelerates *de novo* lipogenesis pathways,^45^ which could offset the effect that decreased digestive exophagy has on lipid droplet accumulation. Together, these results confirm a role for these compounds in digestive exophagy, demonstrating that our screening protocol can effectively identify modulators of lysosome exocytosis during digestive exophagy.

**Figure 4:**
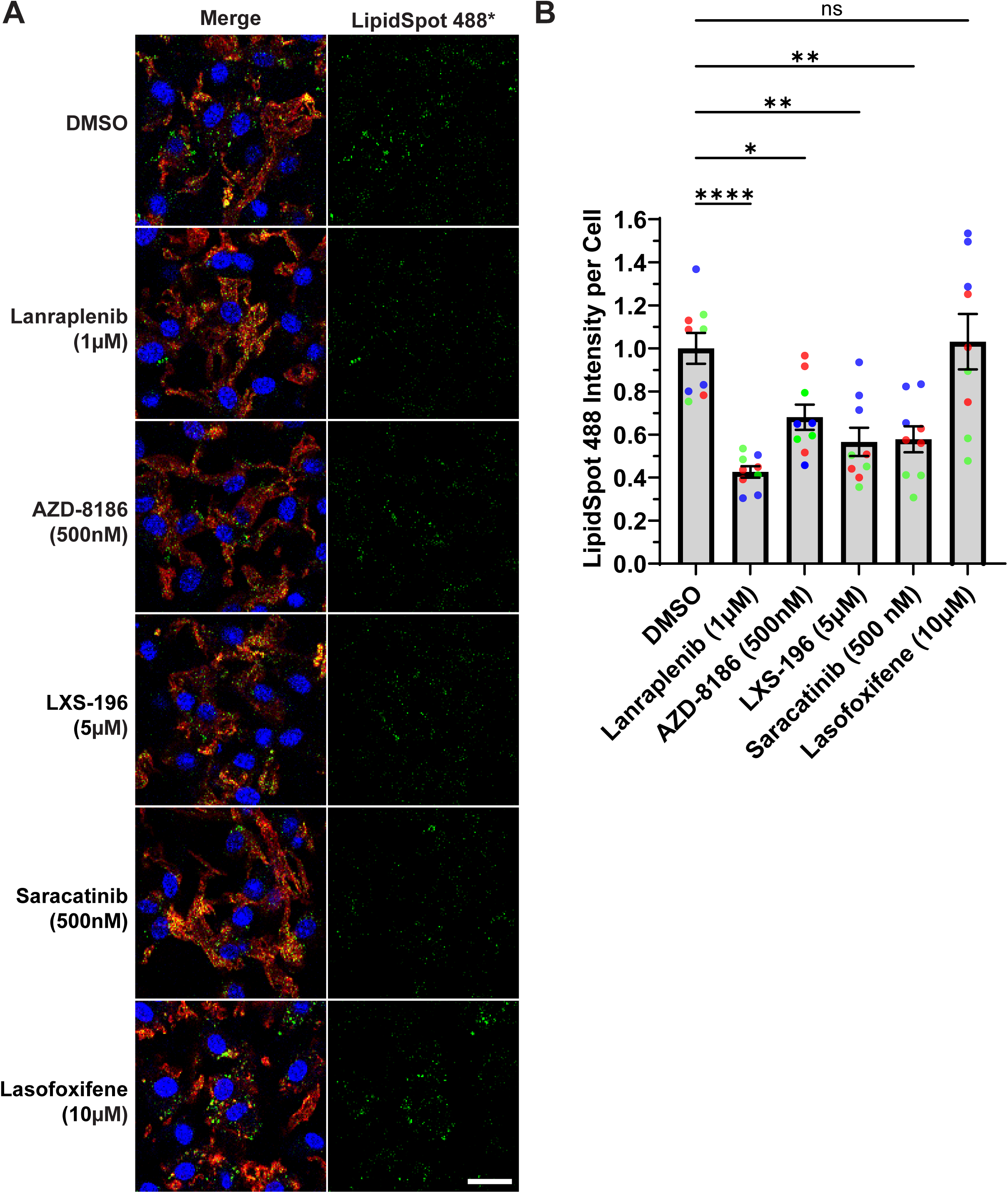
agLDL-mediated foam cell formation is affected by hits from the screen. (**A**) BMMs were treated with DMSO or the indicated drug and assayed for lipid droplet accumulation following a 6-hour incubation with Alexa647-agLDL in a 96-well plate. Maximum projection of LipidSpot 488 signal after the exclusion of agLDL signal (right). Scale bar = 20μm. (**B**) Quantification of the LipidSpot 488 intensity per cell.

### Evaluating whether the five compounds affect oxidized LDL-induced foam cell formation

Because macrophages can interact with modified LDL in both monomeric and aggregated forms in the intima of arteries, we wanted to test whether the compounds we identified in our high-content screen specifically affect macrophage interaction with agLDL or whether they have similar effects on macrophage uptake of modified monomeric LDL. To quantify the accumulation of lipid droplets in macrophages treated with monomeric oxidized LDL (oxLDL), BMMs were treated with oxLDL for 24 hour and stained with LipidSpot 610. We tested how each of the five validated compounds affected oxLDL-induced lipid droplet accumulation in BMMs (Figure 5A, B). AZD-8186, lanraplenib, and saracatinib treatment did not affect oxLDL-induced lipid droplet accumulation compared to a DMSO control. Interestingly, LXS-196-treated BMMs resulted in a decrease in oxLDL-induced lipid droplet biogenesis while lasofoxifene-treated BMMs resulted in an increase in oxLDL-induced lipid droplet biogenesis, again, perhaps due to acceleration of *de novo* lipogenesis pathways.^45^ The result that most of the tested compounds have different effects on lipid droplet accumulation when treated with agLDL compared to oxLDL confirms that the uptake mechanisms for these two LDL species are quite different.^26^

**Figure 5:**
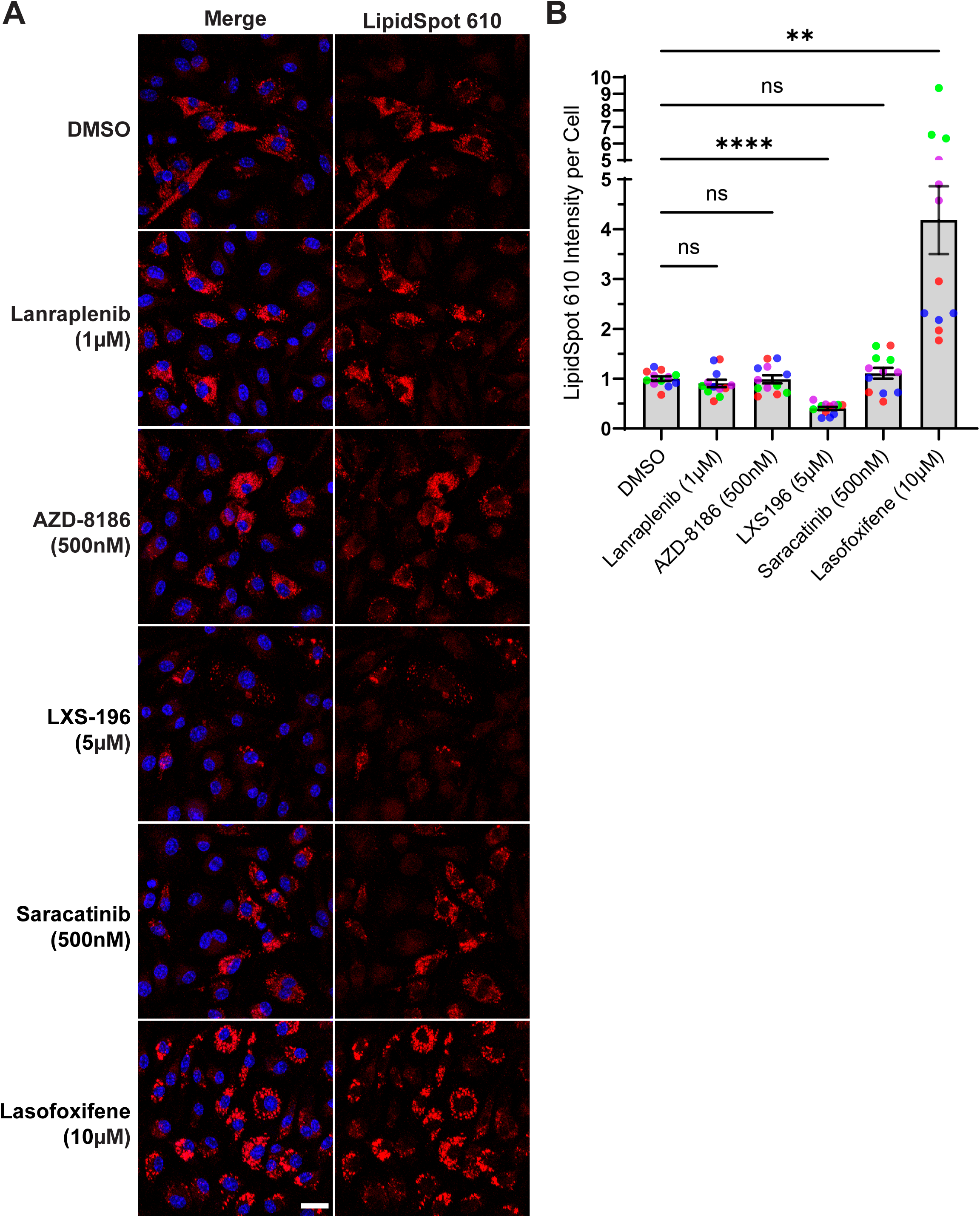
oxLDL-mediated foam cell formation of BMMs treated with hits from the screen. (**A**) BMMs were treated with DMSO or the indicated drug and assayed for foam cell formation following a 24-hour incubation with oxLDL (50μg/ml) in a 96-well plate. Maximum projection of LipidSpot 610 signal (right). Scale bar = 20μm. (**B**) Quantification of the LipidSpot 610 intensity per cell.

## Discussion

During early development of atherosclerosis, macrophages are recruited to the intima of arteries to digest excess modified LDL.^14^ This process causes macrophages to become lipid-laden and is viewed to be a critical event in atherosclerotic progression. Most studies have examined the interaction between macrophages and modified monomeric LDL (often oxLDL), but how macrophages interact with agLDL to become foam cell is less clear.^16,34^ Given that most antioxidant interventions had marginal or no beneficial outcomes,^4^ inhibition of agLDL digestive exophagy could be an alternative therapeutic target for atherosclerosis, especially since most of the LDL in atherosclerotic lesions is aggregated and tightly bound to the extracellular matrix ^17-19^

In this study, we performed a high-content microscopy screen on a repurposed drug library consisting of 2032 compounds to identify novel modulators of digestive exophagy. We identified more than 100 compounds that inhibited lysosome exocytosis more than 60% in J774 immortalized macrophages without severely impairing cell proliferation. We chose to further characterize five of these compounds in BMMs. Three of the five compounds, lanraplenib, AZD-8186, and LXS-196, inhibit targets known to be important for digestive exophagy.^27,36^ The SRC inhibitor saracatinib was the strongest hit from the screen. The estrogen receptor modulator lasofoxifene is approved in the EU to treat osteoporosis in postmenopausal women and has been in clinical trials for osteoporosis treatment in the U.S.^46^ Although lasofoxifene only inhibited lysosome exocytosis by 65%, we decided to further characterize it due to the similarity between osteoclast bone resorption and macrophage digestive exophagy.^29^ All five compounds inhibited lysosome exocytosis in a concentration-dependent manner and reduced F-actin-mediated extracellular compartment formation around agLDL.

Other than lasofoxifene, all compounds impaired agLDL-induced foam cell formation. Estrogen receptor α knockout in mice upregulates lipogenesis in bone marrow cells.^45^ Therefore, it is possible that the reduction of agLDL-induced foam cell formation was masked due to the increase in the basal level of lipogenesis. Consistent with this theory, lasofoxifene treated BMMs showed a strong increase in oxLDL-induced foam cell formation, further suggesting that lasofoxifene specifically inhibited macrophage-agLDL interaction while upregulating *de novo* biogenesis of lipid droplets. Lanraplenib, AZD-8186, and saracatinib did not impact oxLDL-induced foam cell formation, suggesting that these compounds specifically inhibited macrophage-agLDL interactions. On the other hand, the PKC inhibitor LXS-196 reduced both oxLDL- and agLDL-induced foam cell formation. Because PKC controls endocytosis of multiple surface receptors,^47-50^ it is possible that PKC inhibition impaired receptor-mediated endocytosis of oxLDL. Because the uptake mechanisms for these LDL species appears largely different, our high-content screen has the potential to identify distinct and potentially novel atherosclerosis-prevention pathways.

Many of the compounds identified in the screen that decrease digestive exophagy have been shown to alleviate atherosclerosis in various disease models, supporting the idea that slowing digestive exophagy may be a useful therapeutic strategy in slowing the development of atherosclerosis. Through a systems immunology-based drug screening pipeline, saracatinib was identified as a candidate drug to combat inflammation in atherosclerosis.^51^ In *Apoe*^*-/*-^ atherogenic mice, saracatinib reduces atherosclerotic plaque burden by nearly 70% at the highest dose tested.^51^ In this study, we independently identified saracatinib as a strong inhibitor of digestive exophagy. Another compound we tested, lasofoxifene, has been shown to lower the risk of major coronary heart disease events and stroke in postmenopausal women with osteoporosis.^46,52^ Coupled with the result that lasofoxifene treatment decreases digestive exophagy without affecting lipid droplet accumulation in cells treated with agLDL, this raises the intriguing possibility that inhibiting digestive exophagy even without affecting foam cell formation may be beneficial in treating atherosclerosis.

Inhibitors targeting other pathways involved in the macrophage inflammatory response, including JAK-STAT, Aurora A kinase and H1 histamine receptor, were also identified in our screen.^53-55^ Interestingly, treating *Apoe*^*-/*-^ atherogenic mice with the JAK2 inhibitor fedratinib, another strong hit from our screen, reduced atherosclerotic plaque formation.^56^ These results all suggest that the activity of macrophage-mediated digestive exophagy can be modified by tuning the macrophage inflammatory response. This is in line with our previous observation that Toll-like receptor 4 is required for digestive exophagy of agLDL.^27^

The compounds discussed here are only the tip of the iceberg of hits from this screen. As previously mentioned, we identified more than 100 compounds that inhibited lysosome exocytosis more than 60%, any of which could prove to be an effective disease-modifying treatment. Consequently, this list of hit compounds is a valuable resource for atherosclerosis research and merits further investigation in preclinical animal studies of atherosclerosis.

Furthermore, our high-content microscopy-based cell assay has tremendous potential as a drug discovery tool. We have demonstrated proof of concept here on a relatively small library that this platform could be expanded to screens of much larger chemical libraries to identify novel compounds that can modulate digestive exophagy and delay foam cell formation independently of oxLDL interaction.

## Materials and Methods

### Reagents

The following reagents are used in this study: DMEM without L-glutamine (#45000-316, VWR), L-glutamine (#25030081, Thermo Fisher Scientific), Penicillin-Streptomycin (# 15140163, Thermo Fisher Scientific), Fetal bovine serum (#100-106-500, GeminiBio), Copper(II) sulfate pentahydrate (#C7631-250G, Sigma-Aldrich), TBARS Assay Kit (#10009055, Cayman Chemical), 50kDa Amicon Ultra-15 centrifugal filter units (#UFC905024, EMD Millipore), Alexa Fluor 647 NHS ester (Alexa647; #A37573, Thermo Fisher Scientific), FITC NHS ester (#46410, Thermo Fisher Scientific), EZ-Link NHS-Biotin (#20217, Thermo Fisher Scientific), streptavidin protein (#21125, Thermo Fisher Scientific), 10kDa amino dextran (#D1860, Thermo Fisher Scientific), LipidSpot 488 (#70065, Biotium), LipidSpot 610 (#70069, Biotium), Alexa546-phalloidin (#A22283, Thermo Fisher Scientific), Hoechst 33342 (#15547, Cayman Chemical), Phorbol 12-myristate 13-acetate (PMA; # S7791, Selleck Chemicals), LY294002 (#S1105, Selleck Chemicals), LXS-196 (#S6723, Selleck Chemicals), lasofoxifene tartrate (#S9520, Selleck Chemicals), lanraplenib (#S9715, Selleck Chemicals), saracatinib (#S1006, Selleck Chemicals), AZD8186 (#S7694, Selleck Chemicals).

### Lipoproteins and dextran

LDL was isolated from fresh human plasma by preparative ultracentrifugation as previously described.^57^ LDL was fluorescently labeled using Alexa647 NHS ester. Streptavidin-labeled Alexa647-LDL was prepared by labeling LDL with Alexa647 and biotin NHS esters followed by incubation with streptavidin protein. LDL was vortex aggregated for 45 seconds to form agLDL.^24^ Copper-oxidized LDL were prepared as previously described.^58^ In brief, 0.1mg/mL LDL in EDTA-free PBS containing 10μM CuSO_4_ was incubated for 20 hours at 37°C. The oxidation was terminated by adding 0.1mM EDTA and LDL was concentrated to 1mg/mL. Level of lipid peroxidation was measured with TBARS assay kit to ensure a reading >30nmol/mg. Biotin-FITC-dextran was prepared by labeling 10kDa amino dextran with biotin and FITC NHS esters.

### Cell and cell culture

Mouse macrophage cell line J774A.1 was obtained from the American Type Culture Collection (Manassas, VA). J774A.1 cells were maintained in DMEM supplemented with 1% (v/v) penicillin/streptomycin, 3.7g/L sodium bicarbonate, and 10% (v/v) FBS in a humidified atmosphere (5% CO_2_) at 37°C and used at low passage numbers. Bone marrow–derived macrophages (BMMs) were cultured as follows.^26^ Bone marrow was isolated from mice aged 6 to 13 weeks. Sterilized femurs were flushed with cold DMEM, and cells were differentiated for 7 days by culture in DMEM supplemented with 10% (v/v) heat-inactivated FBS, 1% (v/v) penicillin/streptomycin, and 20% L-929 cell conditioned medium in a humidified atmosphere (5% CO_2_) at 37°C.^59^

### Animals

Wild-type C57BL/6J mice (The Jackson Laboratory) were sacrificed for BMMs. Mice were housed in a pathogen-free environment at Weill Cornell Medical College and used in accordance with protocols approved by the Institutional Animal Care and Utilization Committees.

### High-content lysosome exocytosis screen

0.25mg/mL biotin-FITC-dextran was delivered to lysosomes by overnight pinocytic uptake. The next morning, cells were chased for 3 hours in medium to ensure delivery to lysosomes. During the chase, 10μL of media was added to the first 23 columns of 384-well plates (#3764, Corning). In column 24, 10μL of media containing 200μM LY294002 was added to the top 8 wells and 10μL of media containing 4μg/mL PMA was added to the bottom 8 wells. 40nL of a 5mM drug stock or DMSO (column 23) was added to the plates using a Janus Automated Workstation (Perkin-Elmer) at the Rockefeller/Weill Cornell Drug Discovery Resource Center. Following the chase, 8,000 J774 cells in 30μL of media were plated in each well of the 384-well plates using a MultiDrop384 Dispenser (Thermo Fisher Scientific) and allowed to settle for 3 hours. 3μg Streptavidin-Alexa647-agLDL in 5μL was subsequently added to each well for 90 minutes. Following a 6x wash with PBS using an ELx405 Microplate Washer (Bio-Tek Instruments), cells were incubated with 200μM biotin in medium for 15 minutes to bind any unoccupied streptavidin sites before cell permeabilization. Following a 6x wash with PBS, cells were then fixed with 0.5% paraformaldehyde for 15 minutes, washed 6x more in PBS, and permeabilized with 1.3% Triton X-100 for 15 minutes to remove intracellular biotin-FITC-dextran. Finally, cells were washed 6x more in PBS and left in PBS. The next day, images were acquired with an ImageXpress MICRO Automated High-Content Imaging System (Molecular Devices) using a 20x dry objective. 4 frames per well were acquired on three different days.

In most experimental wells, a total of >2,500 cells per condition were imaged and subjected to quantification, though some drug treatments killed cells so fewer cells were imaged in those conditions. To quantify lysosome exocytosis, images were analyzed using MetaXpress (Molecular Devices) image-analysis software. First, all images were corrected for slightly inhomogeneous illumination as described previously.^60^ Briefly, an image was created by averaging all the images from a plate and smoothing the averaged image using a low-pass filter. Then, each pixel in an image was multiplied by the average intensity of the shading image, and the resulting pixel intensities were divided by the shading image on a pixel-by-pixel basis. After illumination correction, background was subtracted from each shading-corrected image by determining the fifth percentile intensity value of the image and subtracting this value from each pixel in the image. Next, a threshold was set in the Alexa647 channel that would include all the fluorescent agLDL in the images.^23,30^ A threshold was also set in the FITC channel that would include all FITC signal in the images. We used the same threshold level for each image within an experimental data set. To remove extraneous signal from fluorescent debris, the “integrated morphometry analysis” module was used to exclude areas of FITC signal larger than 2000 pixels. Total FITC fluorescence intensity within the overlap of the FITC and Alexa647 thresholded areas was measured for each field. The LDL area in each frame was calculated using the same threshold in the Alexa647 channel and counting the pixels that contained LDL signal. Nuclei were counted using the “Count Nuclei” module. Exocytosed biotin-FITC-dextran per field was normalized to both LDL area and cell count. In each well, this measurement is expressed as a percentage of the average of the DMSO controls on its corresponding plate.

### Lysosome exocytosis^23^

1mg/mL biotin-FITC-dextran was delivered to lysosomes by overnight pinocytic uptake. The next morning, cells were chased for 3 hours in medium to ensure delivery to lysosomes. 50μL of medium containing DMSO or drugs at three times the indicated concentration were added to a 96-well plate (#P96-1.5P, Cellvis). Following the chase, 38,000 BMMs in 100μL of full growth media were plated onto each well of the 96-well plate and allowed to settle for 3 hours. Cells were subsequently incubated with Streptavidin-Alexa647-agLDL for 90 minutes. Next, cells were incubated with 200μM biotin in medium for 15 minutes to bind any unoccupied streptavidin sites before cell permeabilization. Cells were then fixed with 0.5% paraformaldehyde for 15 minutes, washed, and permeabilized with 1% Triton X-100 for 10 minutes to remove intracellular biotin-FITC-dextran. Images were acquired with a Leica Stellaris 8 laser scanning confocal microscope using a 20x (0.75 NA) dry objective. 6 frames per well were acquired for two wells per day on three different days.

### Actin polymerization^30^

38,000 macrophages were plated onto each well of a 96-well plate overnight. The next day, cells were pre-treated for 1 hour with DMSO or the indicated drug. To visualize F-actin (filamentous actin), macrophages were incubated with Alexa647-agLDL for 1 hour, washed with PBS, and fixed for 15 minutes with 3% paraformaldehyde. Sufficient Alexa647-agLDL was added to cells to allow Alexa647-agLDL to interact with almost every cell. Cells were subsequently washed with PBS and incubated with 0.02U/mL of Alexa546-phalloidin in 0.05% (w/v) saponin in PBS for 1 hour at room temperature. Images were acquired using a Leica Stellaris 8 laser scanning confocal microscope with a 20x (0.75 NA) dry objective. 6 frames per well were acquired for two or three wells per day on three or more different days.

### agLDL-mediated foam cell formation^28^

38,000 macrophages were plated onto each well of a 96-well plate overnight. Cells were pre-treated for 1 hour with DMSO or the indicated drug, then incubated with Alexa647-agLDL for 5 hours at 37°C and 5% CO_2_. Cells were fixed with 2% paraformaldehyde for 15 minutes and washed with PBS. Cells were stained with LipidSpot 488 (1:1,000) for 30 minutes at room temperature right before imaging. Images were acquired using a Leica Stellaris 8 laser scanning confocal microscope with a 20x (0.75 NA) dry objective. 6 frames per well were acquired for three wells per day on three different days.

### oxLDL-mediated foam cell formation

30,000 macrophages were plated onto each well of a 96-well plate overnight. Cells were pre-treated for 1 hour with DMSO or the indicated drug, then incubated with oxLDL (50μg/ml) for 24 hours at 37°C and 5% CO_2_. Cells were fixed with 2% paraformaldehyde for 15 minutes and washed with PBS. Cells were stained with LipidSpot 610 (1:1,000) for 30 minutes at room temperature right before imaging. Images were acquired using a Leica Stellaris 8 laser scanning confocal microscope with a 20x (0.75 NA) dry objective. 6 frames per well were acquired for three wells per day on three different days.

### Low-content image analysis and quantification

For low-content microscopy experiments, in every experiment, a total of >2,500 cells per condition were imaged and subjected to quantification. For image quantification, MetaMorph software (Molecular Devices) was used. All images subjected to comparative quantification were acquired on the same day using the same microscope settings. For each experiment, we obtained a z-stack encompassing the entirety of the cell for each field at wavelengths appropriate for Alexa647-agLDL, Hoechst, and biotin-FITC-dextran or Alexa546-phalloidin or LipidSpot 488 or LipidSpot 610 as appropriate.

To quantify lysosome exocytosis and actin polymerization, we determined a threshold in the Alexa647 channel that would include all the fluorescent agLDL in the images.^23,30^ After background subtraction in the biotin-FITC-dextran or Alexa546-phalloidin channel, we measured the total fluorescence intensity within the thresholded area for each field. We used the same threshold level for each image within an experimental data set. Nuclei were counted using the “Count Nuclei” module in MetaMorph. Exocytosed biotin-FITC-dextran per field was normalized to cell count in the field. In actin polymerization experiments, total Alexa546-phalloidin staining was also measured in the background subtracted images. Phalloidin staining that colocalized with agLDL was normalized to total phalloidin staining in the field.

Lipid droplet formation was measured using LipidSpot dyes. To quantify agLDL-mediated foam cell formation, we determined a threshold in the Alexa647 channel that would include all the agLDL in the images. Because LipidSpot dyes stain neutral lipids (both agLDL and lipid droplets), this thresholded area was used to exclude LipidSpot 488 fluorescence signal that came from Alexa647-agLDL, so the only signal included came from lipid droplets.^28^ After excluding agLDL signal and background subtraction in the LipidSpot 488 channel, we measured the total fluorescent LipidSpot 488 signal per field. Nuclei were counted using the “Count Nuclei” module in MetaMorph. Data was normalized to cell count. To quantify oxLDL-mediated foam cell formation, we measured the background subtracted total fluorescent signal in the LipidSpot 610 channel and normalized to cell count.

### Statistical analysis

Quantification results from MetaMorph were logged in Microsoft Excel 365 for data processing. Final numbers were imported to GraphPad Prism 9 to be plotted and analyzed. Repeats performed on the same day are indicated in the same color. Error bars indicate SEM unless otherwise noted. Brown-Forsythe and Welch (two-tailed, unequal variance) ANOVA tests with multiple comparisons were performed to test for statistical significance. ns=p>.05, *=p<.05, **=p<.01, ***=p<.001, ****=p<.0001.

## Acknowledgements

This work was funded by grant R01HL093324 from the National Heart, Lung, and Blood Institute (NHLBI). Cheng-I J. Ma (24POST1193887) and Noah Steinfeld (24POST1191688) are supported by postdoctoral fellowships from the American Heart Association. We would like to thank Fraser Glickman and his staff at the Rockefeller/Weill Cornell Fisher Drug Discovery Resource Center for their help developing the screening protocol and carrying out the screen.

We also thank Harold “Skip” Ralph in the Weill Cornell Automated Optical Microscopy Core for his help imaging the 384-well plates from the screen and quantifying lysosome exocytosis from these microscopy images.

## Figure Legends

**Figure S1: Comparing results from different days of the screen**. (**A**-**C**) On all graphs, experimental wells are plotted in black, DMSO controls are in green, LY294002 controls are in magenta, and PMA controls are in orange. A line of best fit is included in blue. (**A**) Results from Day 1 of the screen were plotted against results from Day 2 of the screen with an R^2^ value of 0.6905. (**B**) Results from Day 1 of the screen were plotted against results from Day 3 of the screen with an R^2^ value of 0.6954. (**C**) Results from Day 2 of the screen were plotted against results from Day 3 of the screen with an R^2^ value of 0.6883.

**Figure S2: Actin polymerization around agLDL during digestive exophagy using lower drug concentrations**. (**A**) BMMs were treated with DMSO or the indicated drug and assayed for actin polymerization following a 1-hour incubation with Alexa647-agLDL in a 96-well plate. Scale bar = 50μm. Arrows highlight some areas of actin polymerization around agLDL. (**B**)

Quantification of the colocalization of phalloidin-stained F-actin with agLDL normalized to total F-actin. (**C**) Quantification of total F-actin staining per cell.

