## Supplementary figures and images for "A high-content microscopy drug screening platform for regulators of the extracellular digestion of lipoprotein aggregates by macrophages"

### Supplementary Figure 1

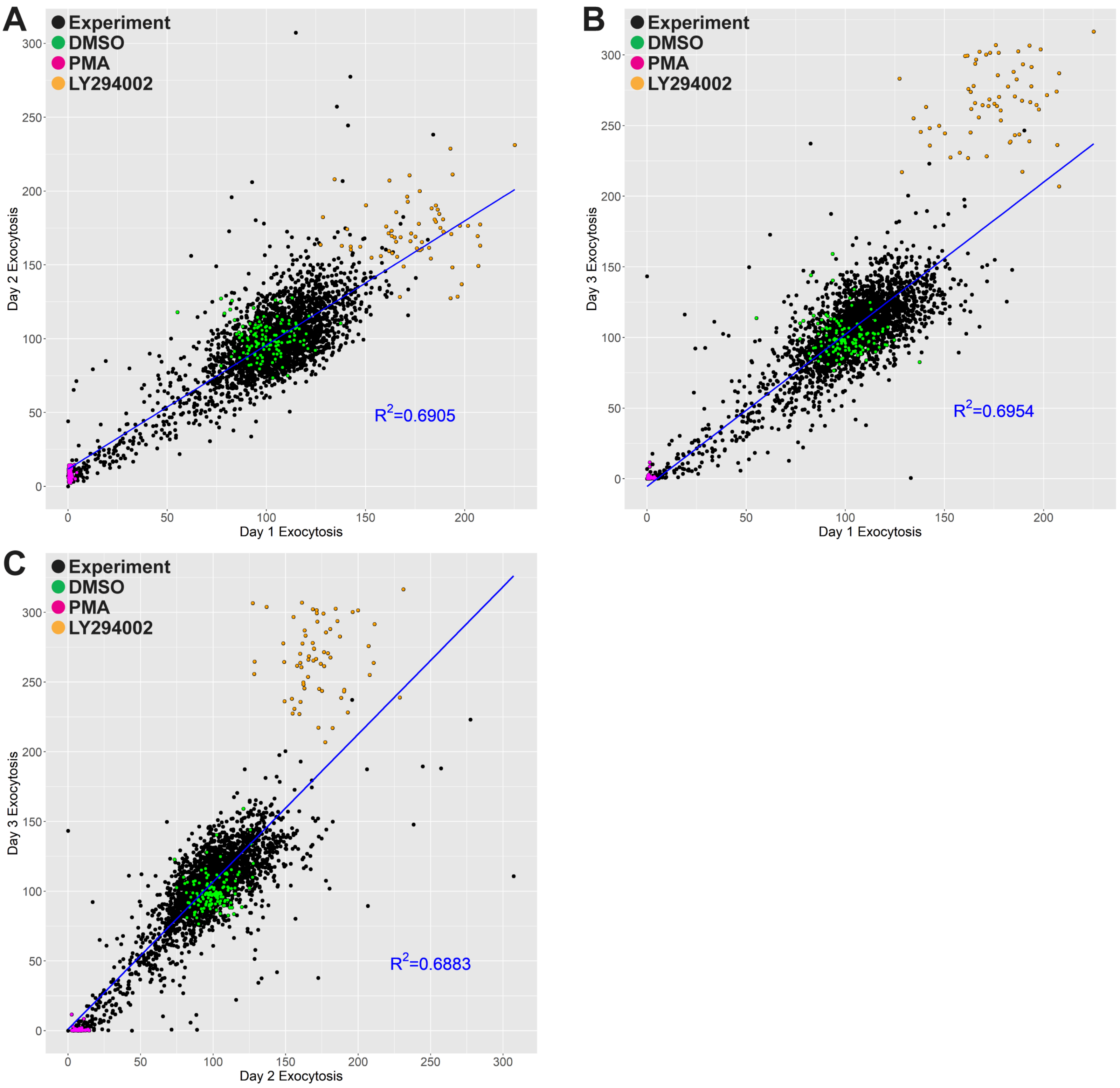

### Supplementary Figure 2

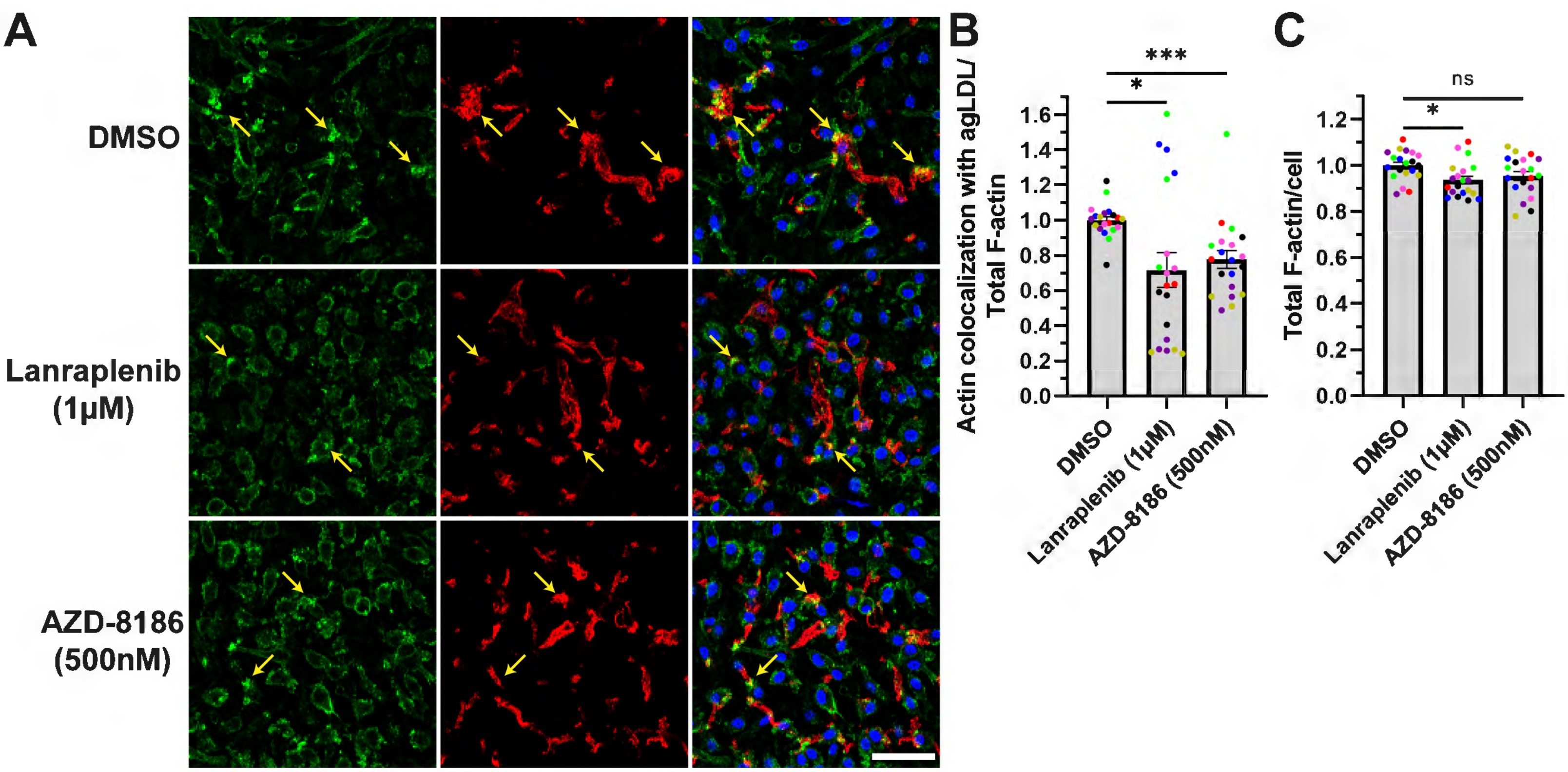
